# Sparse, random sampling is sufficient for central tolerance

**DOI:** 10.64898/2025.12.09.693230

**Authors:** Hannah V. Meyer, Sanjoy Dasgupta, Amitava Banerjee, Yong Lin, Rishvanth K. Prabakar, Sarah R. Chapin, Carl Kingsford, Saket Navlakha

## Abstract

Negative selection in the thymus limits autoimmunity by eliminating T cells that react strongly to self. Individual T cells, however, are only exposed to a small fraction of all self peptides during their “training” in the thymus, and it is puzzling how tolerance can be generalized to the remaining “test” self peptides across peripheral tissues in the body. Using a machine learning perspective, we show that such generalization is possible because the immune system satisfies two conditions: first that peptide abundance levels in the human thymus and periphery are highly correlated (i.e., training distribution ≈ test distribution), and second that cross-reactivity allows T cells to effectively learn binding information of similar peptides without explicitly interacting with all of them. Together, we show that sparse, random sampling of only 10% of self peptides in the thymus is sufficient to avoid reactivity to 90% of peripheral self, and we support this result with diverse experimental data. We then validate two predictions by our model; the first is that only 200–250 antigen presenting cells need to be seen by a T cell to ensure its robust selection, and the second relates how peptides missing from the thymus can drive auto-immunity of peripheral tissues. Overall, we provide a plausible answer to a long-standing question underlying adaptive immunity, and we highlight how generalization, a fundamental challenge faced by nearly every learning algorithm, is uniquely tackled by the immune system.

## Introduction

Negative selection in the thymus [1, 2] is a crucial process for developing a T cell immune response that can eliminate infected or malignant cells in the body while sparing their healthy counterparts. At first glance, however, this process is a numbers game that seems doomed to fail. Developing T cells express receptors (TCRs) on their surface, which are screened for reactivity against self peptides presented by major histocompatibility complexes (MHCs) on thymic antigen presenting cells [3–5]. A T cell that is reactive to any sampled peptide-MHC complex (pMHC) is either deleted or redirected into a regulatory T cell lineage as a safeguard from autoimmune responses. Each developing T cell, however, encounters only a small fraction of the millions of possible self peptides during its short dwell time in the thymus [6–9]. How then do T cells generalize tolerance from the small thymic training sample to the remaining “unseen” self peptides across the body to avoid autoimmune responses?

In reality, this training process is far from perfect; self-reactive T cells escape negative selection and are well-known to exist in the periphery [10–14]. Consequently, there has been intense focus on how peripheral tolerance mechanisms [15] — e.g., regulatory T cells [16], quorum sensing [17, 18], anergy [19] — help to reduce the chance that these self-reactive T cells drive autoimmunity. However, peripheral tolerance alone is not sufficient; the loss or suppression of negative selection can directly result in autoimmunity. The most evident example is the disease Autoimmune Polyglandular Syndrome type 1 (APS1) [20], which is caused by genetic mutations in an important regulator gene of negative selection [21, 22]. This raises the question of how sparse sampling of self peptides during negative selection can possibly generate a T cell population that avoids gross self-reactivity.

To address this challenge, we developed a theoretical model of negative selection that, unlike prior models [9, 17, 23–25], integrates experimental estimates of: (a) peptide abundance levels in the thymus and periphery using whole tissue-level expression data [26, 27]; (b) the number of peptides a developing T cell sees in the thymus using single-cell expression of antigen presenting cells; and (c) T cell cross-reactivity sizes using a state-of-the-art method trained on a large database of mutational scans [28]. Together, we show that central tolerance can be achieved through sparse, random sampling.

## Results

### The immune system satisfies two conditions necessary for generalization

From a machine learning viewpoint, there are two necessary conditions that allow a learning algorithm to generalize to correctly classify data that it has not been trained on [29].

#### Condition 1

The first condition is that the training and testing data must be correlated; i.e., the two must come from the same or a very similar underlying distribution [29, 30]. In our case, the training and testing data correspond to the abundance levels (weights) of self peptides in the thymus and peripheral tissues in the body, respectively. Ideally, a highly abundant peptide in the periphery should also be highly abundant in the thymus, so that any developing T cell that binds such a peptide would, with relatively high probability, encounter the peptide during training and be deleted [31]. Indeed, mismatches between the two distributions — e.g., peptides with high peripheral expression but low thymic expression — are known to be targets of autoimmunity [32–35].

To evaluate condition 1, we derived peptide abundance levels from bulk gene expression data of 29 tissues in the human body (GTEx [27]), as well as of human medullary thymic epithelial cells [26] (Figure 1A–B), which represent the major source of training peptides. To go from gene expression to peptide abundance, we mapped each gene to its corresponding protein using the human reference proteome, and then we mapped each protein to all of the peptides that comprise it using a sliding window. Each peptide’s abundance was taken as the sum of gene expression values of all genes that map to the peptide. We focused on peptide sequences that are 9 amino acids long (9mers), which are the most abundant peptides bound on MHC class I molecules and serve as the sampling space for cytotoxic CD8 T cells. Consequently, we only considered peptides that are visible to these T cells; i.e., peptides that bind to MHCI molecules. We predicted binding to the well-studied HLA-A0201 allele, which yielded 434,276 9mers out of the total 11,136,576 reference proteome peptides. Finally, to home-in on the positions most relevant for T cell binding, we reduced this set of 9mers to 6mers by excluding positions 1, 2, and 9 from each self peptide [24]. Positions 3–8 show large sequence diversity, reflective of the large diversity of TCRs that bind them. In contrast, positions 2 and 9 are tightly constrained because they are typically MHC anchor residues — mutating them prevents peptides from being presented to T cells in the first place [25, 36–38] — and mutations in position 1 tend to have marginal affects on T cell binding [24, 37, 38]. After this reduction, there were a total of 𝑁 = 426,316 self peptides.

**Figure 1.**
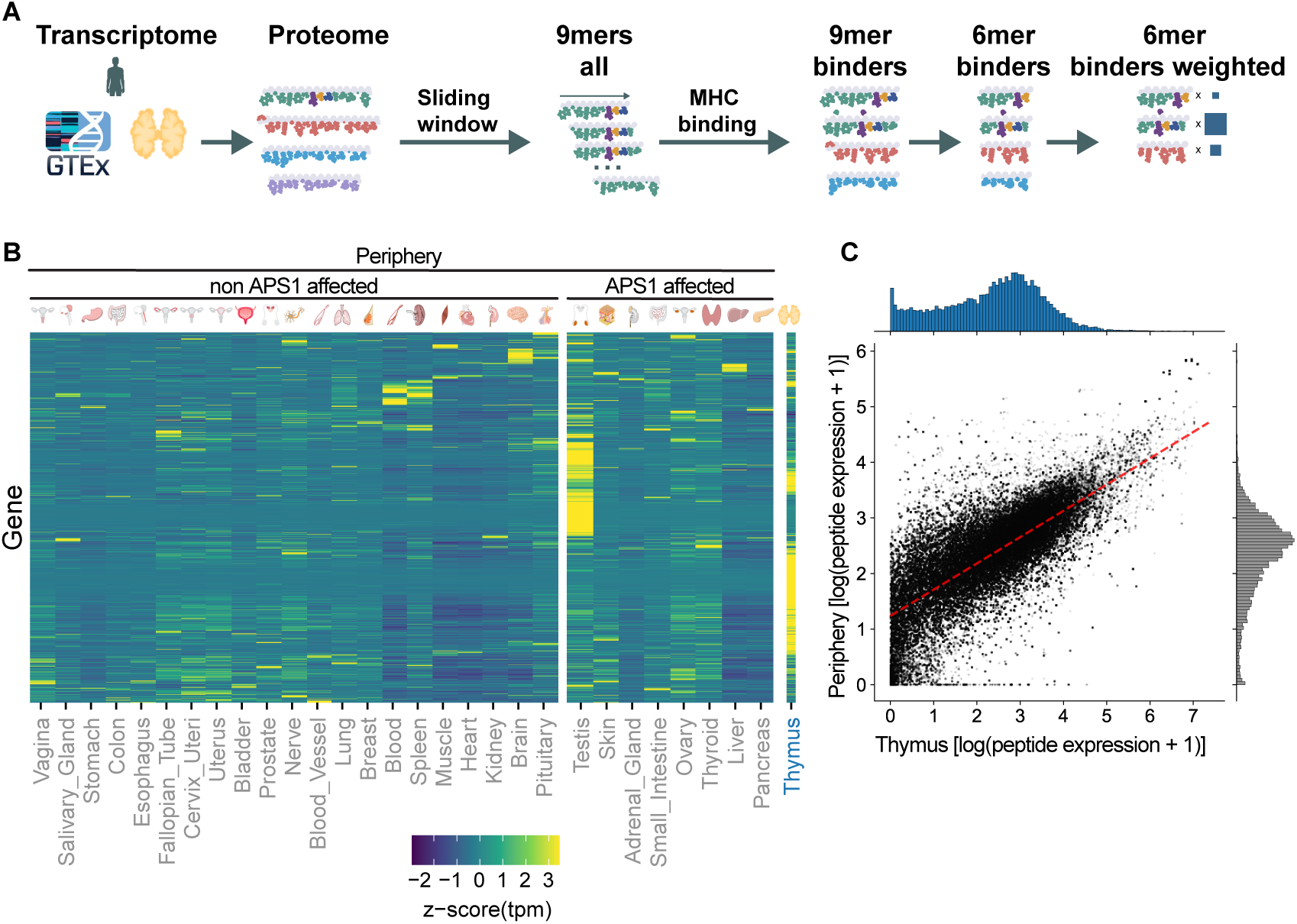
Peptide abundance levels in the thymus and periphery are highly correlated. **(A)** Pipeline to generate MHC-binding weighted 6mer peptides starting from gene expression data. Peptide weights are indicated as blue boxes in the last step of the pipeline. **(B)** Gene expression as z-score of transcript per million (tpm) from 29 tissues in GTEx, as well as thymic epithelial cells [26]. For visualization, the z-score maximum is capped at the absolute value of the minimum, yielding a symmetric scale. Tissues affected and not affected in Autoimmune Polyglandular Syndrome type 1 (APS1) are grouped separately. **(C)** Correlation of normalized thymus (x-axis) and peripheral (y-axis) expression of 6mer MHC-binding peptides, with Pearson’s 𝑟 = 0.76 (dotted red line). For visualization, we plot in log space.

The abundance levels of self peptides in the thymus and periphery were highly correlated (Pearson 𝑟 = 0.76, Spearman 𝜌 = 0.75; Figure 1C). The distributions were also highly non-uniform with abundance levels varying by over 7 orders of magnitude. Moreover, only 0.6% (2494 out of the 426,316) peripheral self peptides were not present (i.e., had zero abundance) in the thymus. This suggests that at the peptide identity level, the thymus contains nearly everything that a T cell could encounter in the periphery (though as we will see, individual T cells only sample a fraction of these peptides during negative selection).

Overall, this means: (a) that under the simplest assumption of random, independent and identically distributed (IID) sampling in the thymus [39, 40], T cells will likely interact with highly abundant peptides; and (b) that these highly abundant peptides are also highly abundant in the periphery, and therefore will likely be protected against. In other words, the “training” ground of the thymus well-approximates the “testing” ground of the periphery, such that interactions experienced in the thymus closely reflect those that will be experienced in the periphery, satisfying the first condition.

#### Condition 2

The second condition is that there must be a notion of similarity between data points such that highly similar points are likely to have similar outcomes. In our case, the data points of interest are pMHC molecules and the outcome of interest is the probability that a TCR binds a given pMHC molecule.

Each TCR can bind many pMHCs (referred to simply as a peptide hereafter), a property called *cross-reactivity* [42–45]. This property means that a T cell not only binds its index peptide (i.e., the peptide that its receptor most strongly recognizes) but also other “similar” peptides at a sufficient strength to trigger an immune response (Figure 2A). We address a TCR based on its index peptide and define the set of peptides it can bind as its cross-reactivity ball [46]. Consequently, a self-reactive T cell (i.e., a T cell with at least one self peptide in its cross-reactivity ball) will escape negative selection if it misses sampling *all* of the self peptides in its ball during negative selection [17]. Equivalently, the T cell will be correctly deleted if it samples *any* self peptide in its ball; crucially, it does not need to encounter most or even many of them. Thus, cross-reactivity provides a mechanism by which T cells can “generalize” [24] information about whether it is self-reactive without having to explicitly interact with its index peptide.

**Figure 2.**
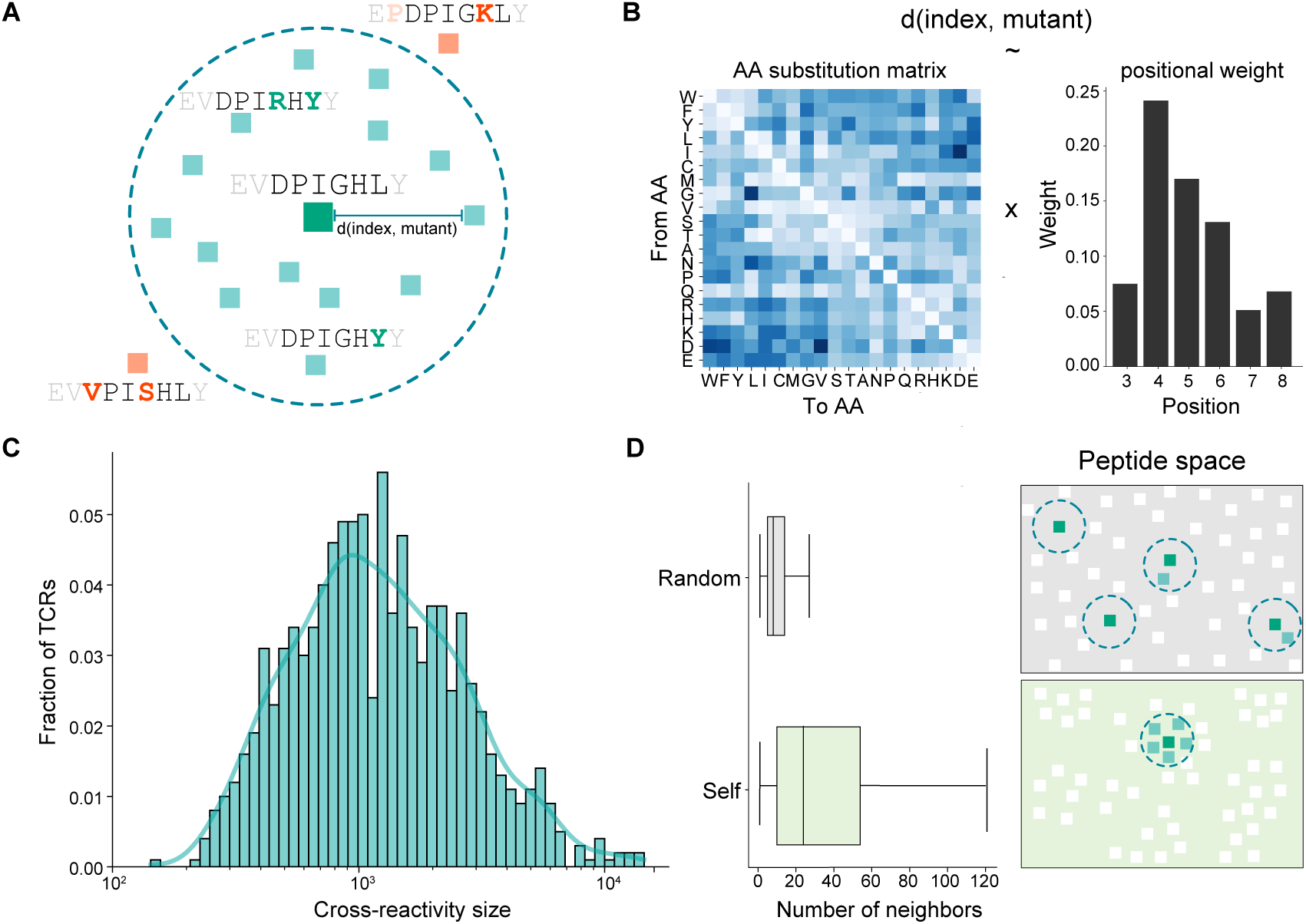
Cross-reactivity provides a mechanism for T cell generalization. **(A)** Each TCR has a cross-reactivity ball surrounding its index peptide (i.e., the peptide that most strongly binds its receptor), indicated by the large green square. Small mutations to the index peptide could preserve binding (peptides inside the circle) or could eliminate binding (outside the circle). Example peptides shown for the a3a TCR [41]. **(B)** The BATMAN cross-reactivity model determines which peptides lie in the cross-reactivity ball of a given TCR’s index peptide. BATMAN computes the distance between the TCR’s index peptide and a given mutant peptide based on an amino acid substitution matrix and a weight on the position of the mutation in the sequence. **(C)** BATMAN generates a broad spectrum of cross-reactivity sizes (i.e., the number of peptides that lie within the ball). Histogram shown for 1000 random TCRs drawn from the space of 64M TCRs. **(D)** If self peptides were randomly distributed in peptide space, there would be on average only 11.9 neighbors in the cross-reactivity ball around a given self peptide. In contrast, for the actual set of human self peptides, there are 44.2 (i.e., 3.7 times more) self neighbors in the cross-reactivity ball around a given human self peptide (left). As a result, the set of human self peptides are more tightly packed in peptide space compared to random peptides (right).

Modeling T cell cross-reactivity requires a notion of similarity between a given TCR’s index peptide and a candidate peptide. We employed a computational method, called BATMAN [28], which recently achieved state-of-the-art performance on a large TCR cross-reactivity prediction benchmark. Given the index peptide of a TCR and a mutated peptide, BATMAN outputs a binding distance of the mutant based on two factors (Figure 2B): a learned amino acid substitution matrix, and a learned weight on each position in the peptide sequence, indicating its importance [38]. These factors allow BATMAN to identify peptides that are similar despite having many mutations separating them. The size of each T cell’s cross-reactivity ball is controlled by a radius cut-off parameter. In accord with prior experimental and theoretical work estimating cross-reactivity sizes (Methods, *Prior work on T cell cross-reactivity*), we set this parameter such that the distribution of sizes has a median of roughly 1000 peptides (from the space of 20^6^ = 64M possible 6mer peptides). Despite using a fixed radius, BATMAN can generate a pre-selection TCR repertoire with cross-reactivity sizes spanning two orders of magnitude, from 100 to 10K peptides (Figure 2C), mimicking the broad spectrum of cross-reactivity sizes found experimentally [23, 47].

The densities of the cross-reactivity balls around each of the 𝑁 TCRs with an index peptide that lies in self reveals the extent to which binding information can be learned from similar self peptides. There were a mean of 44 (median 24) other self peptides in the cross-reactivity ball of a given T cell, compared to a mean of 12 (median 8) if the set of self peptides were instead randomly distributed in the 20^6^ space (Figure 2D). This implies that self peptides are 3–4 times more tightly packed in peptide space than random (Figure 2E), and consequently, that cross-reactivity provides a T cell many more “outs” by which it can learn if it is self-reactive. From a machine learning perspective, the relative compactness of the self distribution means that there is structure in the data that makes generalization more likely to succeed.

In summary, the similarity in the train and test sets (i.e., the abundance levels of peptides in the thymus and the periphery), along with the ability to generalize binding information across a few dozen self peptides via cross-reactivity, together provide two key ingredients for immune generalization.

### A theoretical model of immune generalization

How much potential peripheral damage (self-reactivity) would T cells cause if negative selection did not exist? How much of this damage is reduced by negative selection as a function of the number of peptides an individual T cell sees during training? Here, we develop a theoretical framework to analytically quantify this ratio that accounts for peptide weights (abundances) and T cell cross-reactivity.

Let 𝒰 be the universe of all 20^6^ peptides of length 6, and let 𝒮 ⊂ 𝒰 be the subset of human MHC-binding self peptides; in our case, |𝒮| = 426,316. During development, each T cell generates a random receptor [48] whose index peptide lies in 𝒰. The goal of negative selection is to eliminate any T cell that binds strongly to a peptide in 𝒮.

Each peptide 𝑠 ∈ 𝒮 has two weights, 𝑇 (𝑠) and 𝑃 (𝑠), corresponding to the abundance of 𝑠 in the thymus and periphery, respectively. These abundances are normalized to form a probability distribution; i.e., ^∑^_𝑠∈𝒮_ 𝑇 (𝑠) = 1, and similarly for 𝑃 (𝑠). Consequently, the probability that a T cell “sees” peptide 𝑠 in a given random sampling step is 𝑇 (𝑠).

Each TCR, which is addressed by its 6mer index peptide 𝑡, has a cross-reactivity ball containing all “similar” peptides that can bind the TCR:

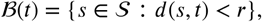

where 𝑑 is the BATMAN distance function with radius 𝑟.

During negative selection, each T cell 𝑡 samples a subset of self peptides, 𝒮^′^ ⊂ 𝒮. If the T cell samples *any* self peptide that lies in its cross-reactivity ball (i.e, if 𝑆^′^ intersects *B*(𝑡)), then the T cell will be deleted. Deletion can only occur for the subset 𝒳 ⊂ 𝒰 of T cells that are self-reactive; that is, for TCRs 𝑡 for which *B*(𝑡) contains at least one self peptide. For our data, 52.8M out of the 20^6^ = 64M total pre-selection TCRs are self-reactive, though the degree of self-reactivity across TCRs varies widely (Figure 2D).

With these definitions, we can compute the probability that a random T cell will cause damage with and without negative selection. Without negative selection, each T cell 𝑡 survives trivially with probability 1: Pr(𝑡 survives) = 1. The damage caused by a random T cell is related to the total peripheral weight of all self peptides that the random T cell can bind:

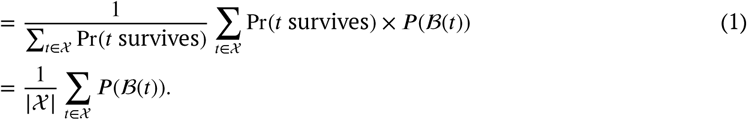

With negative selection, each T cell survives only if it does not interact with *any* self peptide in its cross-reactivity ball after 𝑘 random samples in the thymus with replacement [17, 49]:

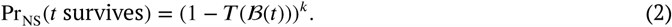

Critically, the probability of survival depends on the thymus weights (𝑇), not the peripheral weights (𝑃). However, damage incurred is calculated using the peripheral weights, since damage occurs in the periphery; hence the importance that the two weights are correlated.

Putting it together, the probability that a random T cell that survives negative selection will cause damage is:

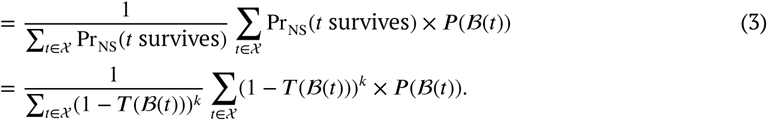

Finally, 1 −Equation (3)∕Equation (1) is the fraction by which negative selection improves protection, which is ideally close to 1.

### How many self peptides does a developing T cell “see” in the thymus?

A critical missing parameter is 𝑘, the number of self peptides sampled by each T cell during negative selection. As 𝑘 increases, the growth rate of the number of unique peptides seen decreases since the same peptide can be seen multiple times (i.e., samples are taken with replacement). This growth rate is further reduced if samples are taken with probability proportional to peptide abundances (Figure 3A).

**Figure 3.**
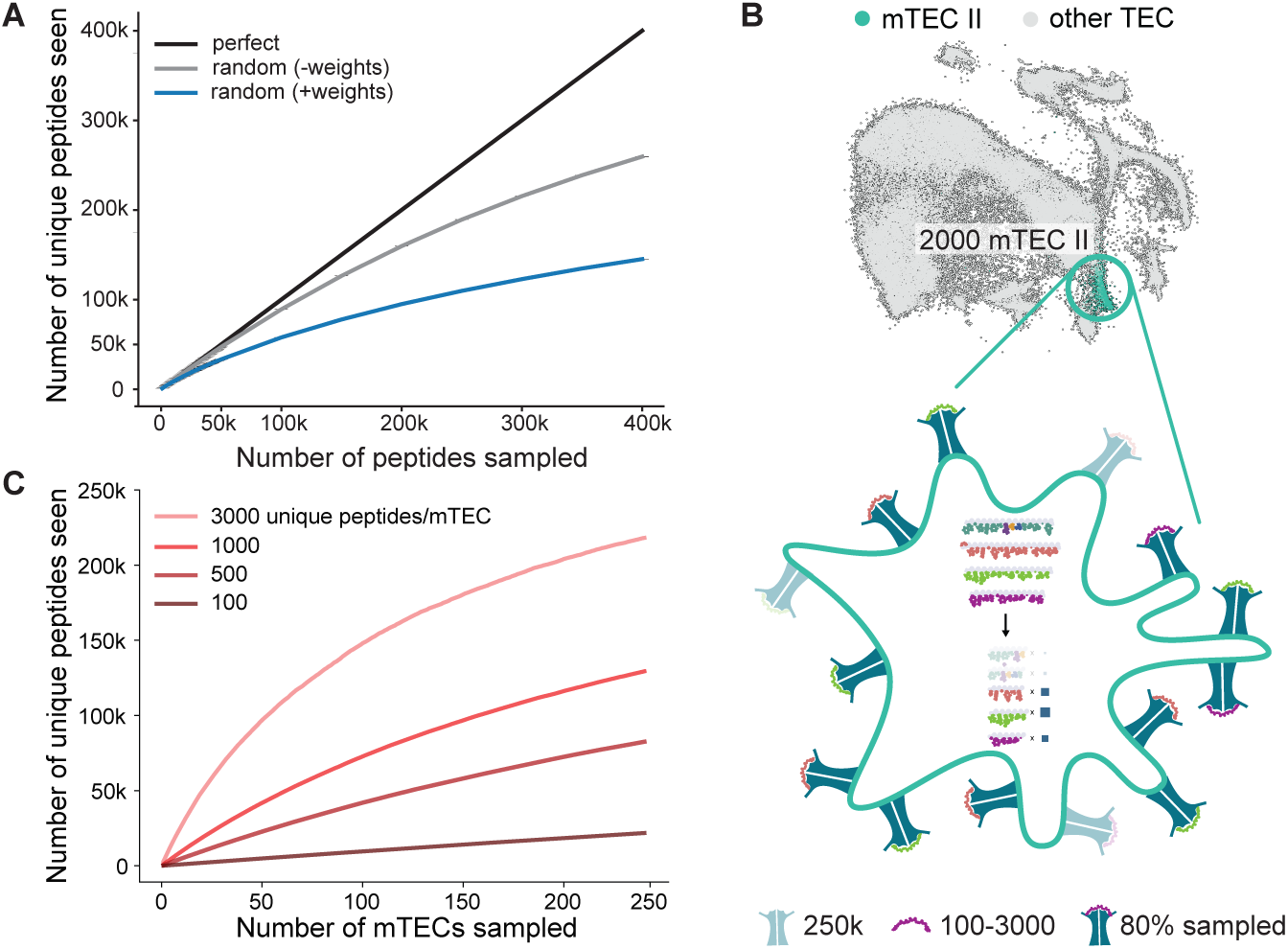
Estimating the number of peptides an individual T cell “sees” in the thymus. **(A)** Relationship between the number of peptides sampled by a T cell and the total number of unique peptides seen. ‘Perfect’ sampling indicates that each sample is unique. ‘Random’ sampling with replacement with uniform weights (‘-weights’) or with peptide abundances (‘+weights’) both show diminishing returns. For example, after 100K samples of peptides (with probability proportional to abundance levels), only 58K unique peptides are seen. **(B)** Pipeline to generate MHC-binding weighted 6mer peptides for medullary thymic epithelial cells with the largest gene expression diversity (mTEC II). Peptide abundances were determined starting from single-cell gene expression data of mTEC II (low dimensional embedding of single-cell data at top of panel) and processed as in Figure 1A (steps summarized inside the highlighted cell in the bottom panel). Peptides for MHC loading were selected based on expression weights (blue boxes) and the number of unique peptide-MHC complexes, which we varied from 100–3000. During simulated scanning of an mTEC, 80% of peptide-MHC molecules are sampled by a T cell (indicated by high transparency level). **(C)** Relationship between the number of mTECs sampled by a T cell and the total number of unique peptides seen. For example, after sampling 240 mTECs with 1000 unique peptides per mTEC, the T cell sees 130K unique peptides.

To provide bounds on 𝑘, we estimated two quantities: the set of self peptides presented across different antigen presenting cells (mTECs), and the number of mTEC interactions by each T cell. For the first quantity, it is difficult to experimentally determine the identities of the self peptides present across the pMHC molecules on the surface of each mTEC. Instead, we estimated this distribution using cell line-derived estimates for MHC abundance [50] and peptide presentation [51–53] as well as single-cell gene expression from 2,000 mTECs of the human thymus (Figure 3B). Based on these, we derived that each mTEC displays between 100 to 3000 unique peptides distributed over roughly 250K pMHC slots on its surface. For each mTEC, we mapped gene expression to peptide abundances; then we sampled 100–3000 unique peptides with probability proportional to its abundance and distributed these peptides proportionally across the 250K MHC slots. When interacting with an mTEC, we estimate that each T cell scans up to 80% of its slots [54], and the union of the peptides across these slots are counted as seen. For the second quantity, we multiplied the average number of mTEC interactions per T cell (5/hour [8]) with the average dwell time in the medulla, focusing on the time with highest susceptibility to apoptosis by negative selection (2 days) [55, 56]. Consequently, each T cell interacts with roughly 240 mTECs. Full details of these derivations are in Methods (*Peptide sampling statistics in the thymus*).

Simulating these 240 mTEC interactions with a range of 100–3000 unique peptides per mTEC, we estimate that each T cell encounters between 20K–200K unique self peptides during its two-day dwell in the thymus (Figure 3C). Thus, T cells are challenged to generalize tolerance to a test set (in the periphery) that is at least as large, or up to 20-times larger, than its training set (in the thymus).

### Sparse thymic sampling is sufficient to protect most of peripheral self

To summarize our model, for each self-reactive TCR 𝑡 ∈ 𝒳, we compute the probability that it samples *any* self peptide in its cross-reactivity ball *B*(𝑡) based on the total weight of the those peptides 𝑇 (*B*(𝑡)) in the thymus (Figure 4A). We then use the total weight of those same peptides 𝑃 (*B*(𝑡)) in the periphery to compute the damage probability (self-reactivity) of the TCR with negative selection (Equation (3)) versus without negative selection (Equation (1)); one minus their ratio tells us how much negative selection reduces damage. Following our estimates of T cell sampling rates, we varied 𝑘 from 30K to 200K, which amounts to roughly 20,000 unique (5% of the total) and 100,000 unique (about 24%) peptides seen, respectively (Figure 4B).

**Figure 4.**
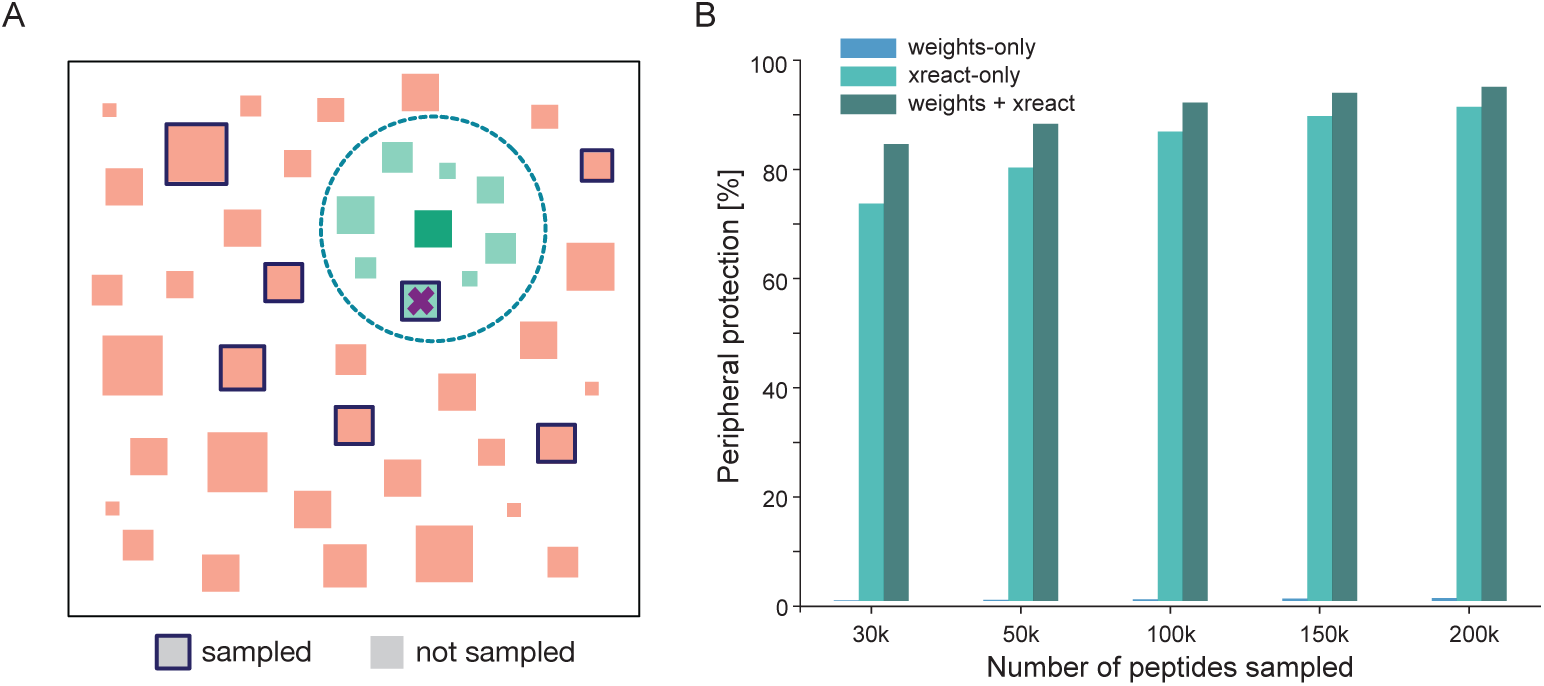
Sparse, random sampling is sufficient to protect most of peripheral self. **(A)** In our theoretical model, individual T cells sample self peptides (outlined squares) according to their abundance (size of square) in the thymus. If the T cell encounters any peptide within its cross-reactivity ball, the T cell is deleted (✖); otherwise, it survives. **(B)** Relationship between the number of peptides each T cell samples and peripheral protection as a result of negative selection. For example, with 100K peptides sampled by each T cell, peripheral protection for the full model (‘weights + xreact’) is 91.3%.

Strikingly, random, sparse sampling by T cells of 5% of unique self peptides (𝑘 = 30K) in the thymus was sufficient to reduce self-reactivity in the periphery by > 80% (Figure 4C). Increasing sampling to 25% of unique self peptides (𝑘 = 200K) reduced peripheral self-reactivity by over 95%. This range, derived under IID sampling assumptions, closely matches experimental estimates of the burden peripheral tolerance faces due to imperfect negative selection (5–20% [10, 13]).

This level of protection required both generalization conditions outlined above. Without correlated peptide weights in the thymus and periphery (for example, by setting all thymus weights to be the same or by randomly shuffling thymus weights), the benefit of negative selection reduces by 5–20% (Figure 4C, xreact-only). Without cross-reactivity, a self-reactive T cell is only deleted if it samples its index peptide, which occurs with near-zero probability (Figure 4C, weights-only). Moreover, our model applied to a random set of 𝑁 self peptides (drawn from the 20^6^ space) offered 15–30% less protection than when applied to the actual set of human self peptides, which further highlights the benefit of cross-reactivity when self peptides lie compactly in peptide space (Figure 2D).

Cross-reactivity was also critical for protecting self peptides with low weight (abundance). The average expression of peptides in the cross-reactivity ball of a high versus a low weight self peptide was very similar; i.e., there was no correlation between the weight of a self peptide and the average weight of other peptides in its cross-reactivity ball (Pearson 𝑟 = 0.09; Supplementary Fig 1). This means that cross-reactivity enables even TCRs with a low-weight self index peptide to be deleted due to having high-weight neighbors.

Overall, these results suggest that negative selection best generalizes to “unseen” peptides using a combination of peptide weights and cross-reactivity. Generalization is further enabled by the relatively compact structure of self peptides in space, and we show is robust across three common human MHC molecules (Supplementary Fig 2 and Supplementary Table 1).

### Model predictions and validation

#### 1. How many mTECs does a T cell need to sample such that its survival probability is invariant to the exact mTECs sampled?

Using the single-cell expression data described above, we grouped mTECs into 𝑚 random, non-overlapping sets, each consisting of 2000/𝑚 mTECs. Each of these sets represents the mTECs sampled by a single T cell during negative selection. For each set, we computed an expression vector corresponding to the sum of expression values for all presented peptides across the 2000/𝑚 mTECs in the set. Using these peptide expression values, we computed the survival probabilities (Equation (2)) for each of the 52.8M self-reactive T cells. Finally, for each pair of sets, we computed the Spearman correlation between the survival probabilities. This analysis includes two parameters — the number of mTECs sampled in each set (which we varied from 20 to 1000) and the number of unique peptides per mTEC (ranging from 100 to 3000). We found that when each set contains 200–250 mTECs, the correlation in the TCR survival probabilities ranged from 84–95% for the two largest estimates for unique peptides per mTEC (Figure 5A). Doubling to 500 mTECs seen per T cell only increases the correlation range to 93–97%, suggesting that doubling development time provides only marginal benefits. This estimate for the number of mTECs sampled by a T cell aligns very closely with our experimentally-derived estimates above (240 mTECs).

**Figure 5.**
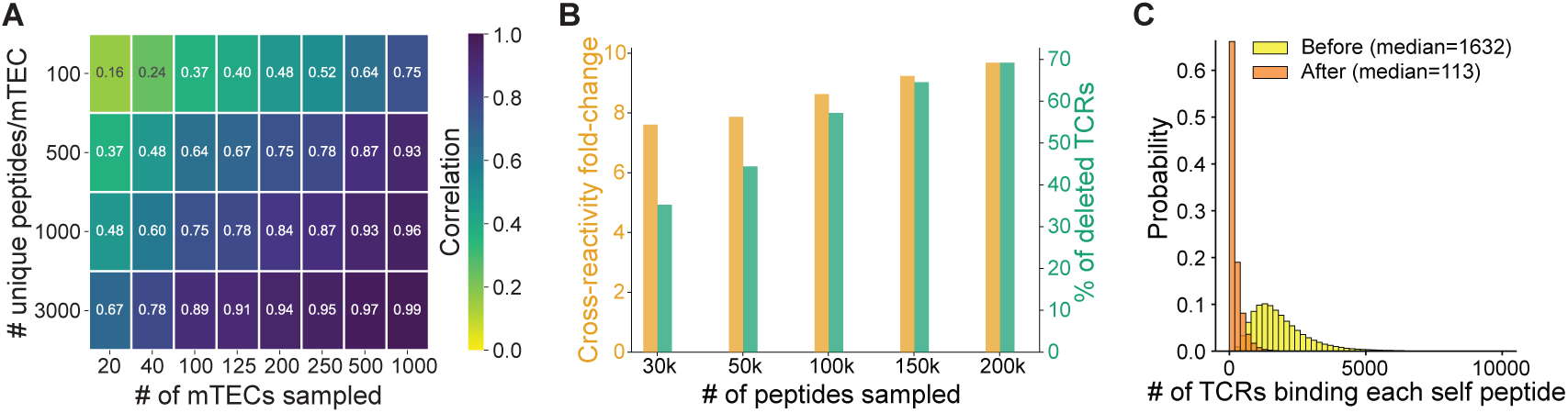
Validating model predictions. **(A)** Heatmap showing the correlation between the survival probabilities of TCRs as a function of the number of mTECs sampled and the number of unique peptides on the surface of the mTEC. For example, with 250 mTECs sampled and 1000 peptides per mTEC, survival probabilities across different mTEC sets visited have a 0.87 Spearman correlation. **(B)** Relationship between the number of peptides sampled by a T cell and the cross-reactivity fold-change of low versus high survival TCRs (left y-axis) or the percentage of deleted TCRs (right y-axis). For example, with 100K peptides sampled, the cross-reactivity size of TCRs likely to survive was 8.6x lower than TCRs unlikely to survive; and the percentage of deleted TCRs was 57.2%. **(C)** After negative selection, the median number of TCRs that recognize each self-peptide is reduced 14-fold compared to the pre-selection repertoire.

#### 2. How well do model estimates on the impact of negative selection on T cell repertoires align with those approximated experimentally?

First, we asked what percentage of the set 𝒳 of 52.8M self-reactive T cells are eliminated by negative selection. In our model, the percentage of deleted T cells can be calculated as:

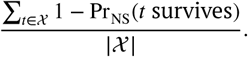

This quantity ranged from 35–70% as a function of 𝑘 (Figure 5B, right y-axis), which is close to the range reported by theoretical and experimental studies (37–75% [49, 57]). Second, we asked how negative selection impacts the range of cross-reactivity sizes of TCRs [58, 59]. We calculated the change in average cross-reactivity towards self peptides as: 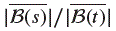, where 𝑠 ranges over all low survival probability TCRs (Pr_NS_(𝑠 survives) < 0.5) and 𝑡 ranges over all high survival probability TCRs (Pr_NS_(𝑡 survives) ≥ 0.5). We found that cross-reactivity to self is narrowed by 7–10-fold comparing low-versus high-likelihood of survival TCRs (Figure 5B, left y-axis). In contrast, cross-reactivity of these TCRs to all peptides in 𝒰 (as opposed to only self peptides) changes by only 2-fold (Supplementary Fig 3), suggesting that the narrowing of cross-reactivity is more targeted towards self-reactive TCRs. Although this distinction between self cross-reactivity and overall cross-reactivity has not to our knowledge been made, these predicted ranges bookend the roughly 5-fold reduction in cross-reactivity after negative selection previously estimated [23, 58, 60].

#### 3. Is our model consistent with alternative mechanisms proposed to overcome the challenge of generalization?

One prevailing theory of peripheral tolerance is quorum sensing [17]. The idea is that a minimum number of TCRs must recognize a self peptide in the periphery to trigger an immune response. As a result, the inevitable leakage by negative selection can be controlled by thresholding responses in the periphery. We found that, before negative selection, each self peptide could bind to a median of 1,632 TCRs from the set of 64M (Figure 5C). After negative selection (using 𝑘 = 200K), this number was reduced to a median of 113 TCRs — a near 15-fold reduction. Thus, quorum sensing in the periphery complements our model and can further reduce the error of generalization and the possibility of autoimmunity.

Thus, our model can recapitulate several important statistics of negative selection and is compatible with a prevailing theory of peripheral tolerance.

### Identifying vulnerabilities in generalization

We asked if our generalization model can identify weaknesses in negative selection that leave certain tissues vulnerable, and if these tissues are implicated in common autoimmune diseases. We first considered all high (> 98%) survival probability TCRs after negative selection and the associated set of self-peptides these TCRs bind. We computed the sum of these peptide abundances in each of the 29 GTEx tissues. We found that six tissues (liver, pancreas, pituitary, salivary, stomach, testis) had between 1.9- and 8.9- fold higher abundance compared to other tissues (Benjamini-Hochberg adjusted p value < 0.05, with empirical p value < 0.01, Supplementary Table 2), and these tissues are implicated in 15 autoimmune diseases (Supplementary Fig 4). This indicates an inherent tolerance vulnerability reflected in tissues implicated in autoimmune diseases.

We next asked if our model could recapitulate the disease phenotype of the Autoimmune Polyglandular Syndrome type 1 (APS1), characterized by mucocutaneous candidiasis, hypoparathyroidism, and adrenal insufficiency. In addition, other tissues implicated in this disease include the liver, skin and pancreas (all APS1 tissues present in GTEx indicated in Figure 1A) [20]. APS1 is caused by genetic mutations in the AutoImmune REgulator (AIRE) gene [21, 22], a crucial transcriptional regulator in thymic epithelial cells. Upon loss of AIRE, as tested in mouse models [3, 61], thymic epithelial cells lose expression of more than 3,000 genes, and mice suffer from subsequent APS1-like autoimmunity as a result of impaired thymic selection.

To test if our model could predict auto-reactivity to tissues afflicted in APS1 as a result of losing AIRE-related peptides, we estimated the effect of AIRE-deletion on thymic expression using human orthologs of murine genes known to be controlled by Aire [3]. Removing these 3,361 genes from human thymus gene expression resulted in 363,344 peptides, i.e., a reduction of 15% compared to the fully intact thymus. We then applied our generalization model using estimated peptides abundances from this impaired thymus; peripheral abundances remain unchanged.

Analogous to our baseline vulnerability analyses above, we considered all high (> 98%) survival probability TCRs in the AIRE-deleted thymus. We found that their associated self peptides were strongly enriched in APS1 tissues (p=1.35E-21, with a 1.6-fold higher abundance; Table 1). The observed level of enrichment was significantly higher compared to when choosing the same number of random TCRs (empirical p-value < 0.01). We also considered all TCRs whose survival probability was 50% higher in the AIRE-deleted thymus compared to the fully intact thymus. The associated peptides of these TCRs were again significantly enriched in APS1 tissues (p=3.89E-04; Table 1), and this level of enrichment was roughly two orders of magnitude higher compared to when choosing random TCRs (empirical p-value < 0.01).

**Table 1.** Generalization model under thymic AIRE deletion predicts peptide enrichment in APS1 tissues. Mean peptide abundance comparison in APS1 versus non-APS1 tissues and their associated p-values (one-sided [>] paired). ‘𝑁_TCRs_’ and ‘𝑁_peptides_’ specify the number of TCRs and their associated target peptides in the ‘model’. For comparison, the first row shows all self-peptides stratified into APS1 affected tissues and non-APS1 affected tissues, with no significant difference in mean abundance.

| model | $N_{\text{TCRs}}$ | $N_{\text{peptides}}$ | p value | mean abundance | |
| --- | --- | --- | --- | --- | --- |
|  |  |  |  | APS1 | non-APS1 |
| all peptides | — | 426,316 | 0.83 | 49.17 | 49.42 |
| > 98% survival ΔAIRE model | 2,803,613 | 82,565 | 1.35E-21 | 18.23 | 11.46 |
| > 50% increased survival ΔAIRE model | 118,464 | 61,515 | 3.89E-04 | 20.39 | 16.99 |

Overall, these results suggest that (a) TCRs that likely survive negative selection preferentially target peptides in tissues linked to autoimmune disorders; and (b) TCRs that likely survive after thymic deletion of AIRE preferentially target APS1 tissues; i.e., APS1 tissues are relatively less protected from autoimmunity compared to non-APS1 tissues. The latter result further highlights the importance of correlated peptide abundances in the thymus and periphery towards accurate generalization.

## Discussion

In biology, the generalization problem is most commonly studied in the brain, where learning a concept from a few examples (e.g., recognizing someone’s face) is a regular challenge. Here, we explored how generalization is also faced during the development of T cells. There are two conditions necessary for any learning system, be it biological or otherwise, to properly generalize. We showed that these two conditions — namely, that the training and testing data are correlated, and similar data points have similar outcomes — are satisfied by the immune system. The former relates to peptide abundance levels in the thymus (training) and periphery (testing), respectively, which to our knowledge have not been considered in previous models of negative selection, and which we showed are highly correlated. The latter relates to the fact that an individual T cell can react to many similar peptides (called cross-reactivity), which serves as a mechanism by which a T cell can learn if it is self-reactive without having to “see” every self peptide. Together, we showed that sparse, random sampling of self peptides in the thymus is sufficient to avoid reactivity to most of peripheral self and that this observations holds across multiple HLA alleles. Finally, we showed that our generalization model can predict vulnerabilities inherent to central tolerance, reflected in enrichment of autoimmune target tissues under both baseline and AIRE deletion conditions.

While we focused on overcoming the challenge of sparse peptide sampling during thymic selection, there may also be some benefits to this strategy. For example, sparse sampling allows T cells to develop faster than complete sampling of all self peptides, which takes exponentially longer since the rate of seeing new peptides diminishes over time. Second, complete sampling would produce a perfectly self-tolerant T cell repertoire but with a deterministic set of “holes” outside of self peptide space, which pathogens could evolve to exploit [10, 13]. In contrast, sparse sampling means that self-tolerance is compromised for smaller, stochastic holes in nonself space. Third, low levels of self-reactive T cells in the periphery may be beneficial for diverse processes, such as wound healing, tissue homeostasis, and T regulatory cell sustenance (as reviewed by Richards *et al.* (2016) [13]). Thus, sparse sampling coupled with peripheral tolerance could be an effective strategy to trade-off generalization amongst speed, discrimination, and other physiological functions.

Our work opens the door to many future questions. First, while our assumption of random, IID samples in the thymus makes our model simple and analytically tractable, T cell encounters may be more structured and encompass a wide spectrum of antigen-presenting cells in the thymus. We focused on the most transcriptionally diverse subset of mTECs; however, T cells also encounter other antigen presenting cells, including mTEC subsets with different transcriptional activity [62, 63], as well as B cells and dendritic cells, which can present both intrinsic antigens and antigens derived by transfer from mTECs [64–66]. Single-cell expression data from these antigen-presenting cells [62, 67] may enable a more comprehensive model of thymus encounters, though quantitative experimental measurements of antigen transfer remain lacking. Second, while our analysis of single-cell mTEC expression allowed us to estimate how many peptides are seen by individual T cells in the thymus, this approach required some assumptions, such as extrapolating the actual set of peptides present on the surface of mTECs from whole-cell expression and cell-line-derived protein and peptide abundance estimates. Although peptide presentation is correlated with both RNA and protein expression levels [68–70], future work performing immuno-peptidomics on antigen-presenting cells would allow for more precise estimates of peptide sampling. Third, while our model of cross-reactivity (Methods, *Prior work on T cell cross-reactivity*) takes into account two factors — biochemical similarity of amino acid substitutions and a weight on each position of the peptide sequence — both of these factors were derived from peptide mutations that only systematically spanned the 1D Hamming distance [28]. Understanding the underlying distribution from which these factors are drawn across TCRs, alongside more expansive mutational scan data, would enable more accurate modeling of the diversity of TCR cross-reactivity sizes. Further, while our focus on 6mer peptides allowed us to explore the 20^6^ = 64M TCR space without sub-sampling, exploring TCR-pMHC interactions in the 9mer space (20^9^ = 5 × 10^11^), while unlikely to be possible exhaustively, could better account for potential allosteric interactions between MHC binding residues and TCR-pMHC binding [71, 72]. Fourth, our model can be used to make HLA-specific predictions of target auto-immune epitopes to detect individuals at risk for tissue-specific autoimmune diseases. Fifth, generalization to avoid self must be balanced against over-generalization, which may hinder detection of nonself [9], given the similarity between self and nonself peptides [24, 73]. Understanding how this fine-line can be achieved is reminiscent of anomaly detection algorithms [74].

Furthering this connection with machine learning, the T cell selection problem includes some unique twists on standard generalization problems. For example, in typical machine learning setups, the training and test sets would not overlap, but in our case, they almost exactly overlap (i.e., the thymus and periphery contain essentially the same set of peptides), although with slightly shifted weights (peptide abundances). This represents a unique type of domain adaptation problem [75], where the training and test distributions are close but not identical (i.e., mild distribution shift). In addition, the training set is typically fully accessible to the model, but in our case, the training set is only partially accessible to each T cell. Versions of this setup are used in instances where each learner (T cell) is trained on a different slice of the training data; for example, in ensemble learning this idea is used to improve robustness, or more recently, in federated learning [76], to preserve privacy. Expansion of these classically studied machine learning paradigms to better model immunological realities could be mutually beneficial to both fields.

## Methods

### MHC and haplotype selection

We analyzed alleles and haplotype frequency data from http://www.allelefrequencies.net [77]. Machine-readable frequency files were kindly provided by Faviel Gonzalez-Galarza. Population ancestries from http://www.allelefrequencies.net/datasets.asp#tag_5 were curated into six broadly continental and one mixed ancestry: Amerindian, Asian, European, Hispanic, Oriental, Sub-Saharan and Mixed. The most common allele is HLA-A0201, which we focused on in our main analyses. To extend our analyses beyond a single HLA-A allele, we chose a haplotype common in 6 out of the 7 ancestries (Supplementary Fig 2): HLA-A0101, HLA-B0801, and HLA-C0701.

### Datasets

Our generalization model relies on peptide abundance estimates from peripheral tissues and thymic antigen presenting cells. We infer the peptide abundances from gene expression data as described in the following sections. For direct comparison of peripheral and thymic expression, we analyzed bulk expression data because consistent bulk technologies were used across a large number of peripheral tissues [27] and from our published thymus study [26]. For estimating sampling statistics and peptide coverage in the thymus, we needed more fine-grained data, and hence used single-cell expression data from human thymic epithelial cells.

#### Bulk RNA sequencing data from 29 human tissues in the Genotype Tissue Expression dataset

We downloaded bulk gene expression from the Genotype Tissue Expression (GTEx) portal (version 8, https://gtexportal.org/home/downloads/adult-gtex/bulk_tissue_expression, accessed June 11, 2024, gene expression TPM from RNASeQCv1.1.9). Using sample mapping (GTEx_Analysis_v8_Annotations_SampleAttributesDS.txt) and TPM files (GTEx_Analysis_2017-06-05_v8_RNASeQCv1.1.9_gene_tpm.gct.gz), we computed the mean expression per gene and per tissue across biological replicates. The final dataset contained 56,200 genes across the following 29 tissues: adrenal gland, bladder, blood, blood vessel, brain, breast, cervix uteri, colon, esophagus, fallopian tube, heart, kidney, liver, lung, muscle, nerve, ovary, pancreas, pituitary, prostate, salivary gland, skin, small intestine, spleen, stomach, testis, thyroid, uterus and vagina.

#### Thymus bulk RNA sequencing data

We used the Transcript per Million (TPM) normalized expression data from immature and mature human bulk-sorted medullary thymic epithelial cells (mTECs) generated by Carter *et al.* (2022) [26] (GEO accession: GSE201719) to obtain mTEC (thymus) peptide weights. We first obtained the mean TPM expression for each transcript across three biological replicates of immature and mature mTEC samples and then summed these TPMs to obtain the total mTEC transcript expression. We then summed all transcripts mapping to a single gene to obtain TPMs for 40,481 genes.

#### Thymus single-cell RNA sequencing data

We re-processed five public single-cell RNA sequencing datasets comprising 23 human thymus samples [62, 63, 67, 78] and seven new single-cell and single-nucleus RNAseq datasets of human pediatric thymus samples through a common pipeline: 1) alignment with STARsolo [79] (reference genome: GRCh38 (primary assembly); Gencode annotation version V44), which includes gene count estimation with expectation maximization and cell filtering to retain singlets; and 2) quality control with *scanpy* v_1.9.3 [80], where cells with mitochondrial gene percentages greater than 15% and with fewer than 200 expressed genes were removed from downstream analysis. We integrated single-cells across studies using scVI v_1.0.3 [81], correcting for sequencing protocol, batch and sample. Then, we identified all epithelial cells in the datasets, followed by iterative subclustering of these cells to obtain 17 thymic epithelial cell types (data analyses manuscript in preparation). For all downstream analyses, we focused on cells that we annotated as mTECII, which are characterized by high AIRE and promiscuous gene expression [26], thus providing an upper bound for gene diversity within a single cell. The low dimensional representation of each cell’s gene expression is computed as latent factors by ‘single-cell annotation using variational inference’ [80] and is shown in Figure 3B, upper panel.

### Peptide abundance

We downloaded the human reference proteome (id: UP000005640, Uniprot release: 2022_02, downloaded July 14, 2022) and, following prior work [24, 73], used a sliding window to generate all possible 9mer peptide sequences. We obtained the unique set of these peptides (11,136,576) and predicted their binding to HLA-A0201 using NetMHCpan 4.1 [82]. We combined weak and strong binders (netMHCpan binding affinity rank < 2)to obtain the set of 434,276 human 9mer peptides with predicted HLA-A0201 binding. For each binder, we removed the 1^st^, 2^nd^ and 9^th^ amino acid to yield 426,316 6mer peptides with predicted HLA-A0201 binding. We mapped each peptide to the protein it was derived from and then to the gene encoding the protein, and assigned each peptide a thymus and peripheral abundance (weight) based on the gene expression values in the bulk thymus and peripheral datasets, respectively. Specifically, we used Ensembl Biomart (version: grcg38.p14, genes110) to extract all ensembl gene ids and corresponding uniprot protein ids; any gene id without corresponding uniprot entry was removed from downstream analyses. Gene identifiers in the thymus (both bulk and single-cell) and peripheral datasets were then mapped to these uniprot ids and each peptide mapping to a given gene was assigned this gene’s TPM expression level. Finally, we calculated unique peptide abundance levels by summing all TPMs for a given peptide.

### Peptide sampling statistics in the thymus

There are four critical parameters needed to estimate the number of peptides sampled per T cell during negative selection: the total number of MHC complexes on the mTEC surface, the number of unique peptides that occupy these complexes, the percentage of pMHC complexes scanned by a T cell per mTEC encounter, and the total number of TCR-mTEC interactions.

- **Total number of MHC complexes on the mTEC surface:** the number of MHC molecules on the surface of an mTEC differs by cell type and can change over time based on infection status and environment. However, across both murine and human unstimulated cells, different MHC classes, and different MHC alleles, roughly 10^5^ molecules have been observed in measurements by immunoprecipitation using MHC-specific antibodies [50, 83, 84]. Here, we chose an upper bound estimate of 250K MHC complexes/cell based on immunoprecipitation using the monoclonal antibody B22 recongnizing the murine MHC I molecule D^b^ on EL4 thymoma cells [50].
- **Total number of unique peptides per mTEC:** a common experimental strategy to estimate the number of unique peptides bound on MHC molecules is to take cells of interest and wash peptides off of MHC alleles by mild acid elution followed by peptide separation by microcapillary high-performance liquid chromatography. After identification of peptides by mass-spectrometry, peptide abundances can be estimated by comparison to a standard abundance curve of synthetic peptides. However, for the above processing pipeline, a starting cell number of 10^8^ is often required. Despite recent progress in reducing the cell numbers down to 10^6^ [85, 86], this assessment remains challenging for rare cell types from limited patient material, such as TECs, and to date there are no comprehensive estimates in these cells. Thus, we extrapolate from data derived from cell lines expressing the HLA-A0201 allele, which detected a range of 100–3000 unique peptides per cell [52, 87]. This estimate is also conserved across other MHC alleles and species [51, 53], allowing us to apply these estimates to other MHC loci.
- **Percentage of pMHC scanned per T cell encounter:** quantitative measurements of pMHC encounters on antigen-presenting cells (APC) such as mTECs are often focused on estimating the lower bound of specific pMHCs required to stimulate a T cell response [88–90]. Here, we are interested not only in response-eliciting pMHC encounters, but an overall estimate of sampling of the pMHC space per APC. We approached this by estimating the surface area explored by a developing T cell upon encounter of an APC in the thymus. We relied on imaging tracks recorded by Bousso *et al* [54, Figure 2], measuring the crawling of fluorescently labeled thymocytes on the APC surface in re-aggregate thymic organ cultures. While variable from encounter to encounter, we estimate a lower and upper bound of surface area covered to be 10–80%. In our analyses, we used the upper bound estimate (80%), though this parameter had the smallest effect on the number of unique peptides seen because each peptide reoccurs many times across pMHC complexes on a cell (i.e., at least 250K/3000 ≈ 83 times).
- **Total number of TCR-APC interactions during negative selection:** the estimated number of interactions between thymocytes and APCs are also derived from thymocyte imaging in a thymus culture system, here, agarose-embedded sections of thymic tissue explants [8]. In addition, traceable thymocytes were introduced in this system by hematopoietic chimerism with fluorochrome-expressing bone marrow. Measuring thymocyte motility and interaction in this system, Le Borgne *et al.* [8] estimated on average 5 interactions between thymocytes and medullary APCs/hour. To estimate the total number of interactions, we consulted estimates from GFP-reporter mice where the level of GFP can indicate how much time has passed since being licensed for negative selection [55]. These experiments conclude that thymocytes spend up to 4 days in the medulla, but are only in an apoptosis susceptible state, i.e., actively undergoing negative selection, for about two days [56]. Together, we estimate about 5 interactions/hour × 48 hours = 240 interactions with APCs for each T cell.

### Prior work on T cell cross-reactivity

Modeling T cell cross-reactivity requires a distance function that can be used to predict all the peptides that activate a given TCR. Extensive progress has been made on the opposite problem — predicting all the TCR sequences that can bind a given peptide [91] — but fewer studies have attempted to characterize the cross-reactivity function itself [92, 93], largely due to insufficient data detailing how small mutations in an index peptide affects TCR binding. Consequently, prior work [25] has modeled cross-reactivity using general sequence-based distance functions, such as r-contiguous matching [24, 74], Hamming distance [73, 94, 95], and BLOSUM distance [38, 96–99], which takes biochemical similarity of substitutions into account.

Determining the size of the cross-reactivity ball has been of intense interest in the literature [43], given its wide-spread importance to pathogen detection, autoimmunity, and transplant rejection [47]. Since it remains daunting to quantify this number comprehensively, most estimates in the literature combine sparse experimental probing of antigenic space with theoretical extrapolations. The reported estimates are also sensitive to the length of the peptide tested and whether MHC binding is considered. For example, Mason (1998) [42] estimated that a single T cell can cross-react to 1M peptides (length 9) without factoring MHC binding, and only 0.1% [100] to 1% [101] of peptides are believed to bind to a given MHC. Consequently, the cross-reactivity size for MHC-binding peptides may be closer to 1–10K. Hybrid experimental-theoretical estimates by Woolridge *et al.* (2012) [101] and Hiemstra *et al.* (1999) [102] support the 1M number and for MHC-binders, but these studies used longer peptides (lengths 10 and 11, respectively), which occupy a 20–400x larger space, and thus again, the average cross-reactivity size for length 9 MHC-binders may be in the low to mid thousands. Wortel *et al.* (2020) [24] considered length 6 peptides, noting that positions 2 and 9 are MHC anchor residues [24, 25, 37, 38] and position 1 mutations have the lowest affect on TCR binding [24, 37, 38]. Within the 6mer space, Wortel *et al.* (2020) estimate that each TCR can bind to one in every 30K peptides [103], which again amounts to a cross-reactivity size of a few thousand.

### Modeling T cell cross-reactivity using BATMAN

To model T cell cross-reactivity, we used BATMAN [28], a recent method employing a hierarchical Bayesian model to predict TCR activation by a given peptide based on its distance to the TCR’s index peptide. This peptide-to-index distance is a product of two factors: (1) a learned amino acid (AA) substitution distance matrix from the index AA to the mutant AA, and (2) a learned weight on each position in the peptide sequence. In the original work, these factors were learned from a large database, called BATCAVE. BATCAVE contains the largest database of peptide mutational scan assays collected to date, covering over 22,000 TCR-pMHC pairs, and 151 mouse and human TCRs tested against 25 mutagenized index peptides.

In our main analysis, we focused on one HLA allele (A02:01) and on 6mer peptides (positions 3–8 of each peptide). Consequently, we re-trained BATMAN using a subset of the BATCAVE data (i.e., only HLA-A*02:01-binding peptides, as predicted by NetMHCpan4.1 [82], resulting in 1,429 TCR-pMHC pairs, containing 10 TCRs binding to 10 unique index peptides). Since the original 9mer BATCAVE data had all peptide positions mutagenized, extracting positions 3–8 could result in multiple identical 6mer peptide entries with potentially different TCR activation levels, even for the same TCR. Nevertheless, in five-fold cross-validation tests for TCR activation prediction, the AUCs were very consistent when using 9mers (AUC=0.709) versus 6mers (AUC=0.704). Moreover, the AA substitution matrices inferred using 9mers versus 6mers were highly correlated (Pearson 𝑅 > 0.99), and, when using all 9 peptide positions, the inferred positional weights of positions 1, 2, and 9 were the smallest compared to positions 3–8. Together, these results further support our focus on peptide positions 3–8 towards accurate T cell cross-reactivity modeling.

For the other individual alleles (A0101, B0801, C0701), as well as the haplotype analysis on the combined alleles (A0101-B0801-C0701), we re-trained BATMAN on the full BATCAVE database, since there was not sufficient data for an allele-specific model.

### Immune disease statistics

Our immune disease statistics (summarized in Supplementary Fig 4) were based on a comprehensive literature survey by Hayter and Cook [104]. They identified 81 autoimmune diseases that fulfill at least two of the following five criteria (as defined by the original study): (1) the specific adaptive immune response is directed to the affected organ or tissue; (2) autoreactive T cells and/or autoantibodies are present in the affected organ or tissue; (3) autoreactive T cells and/or autoantibodies can transfer the disease to healthy individuals or animals; (4) immunization with the autoantigen induces the disease in animal models; and (5) elimination or suppression of the autoimmune response prevents disease progression or even ameliorates the clinical manifestation. We referred to Table 2 in the original publication, matching disease name (as indicated) with target tissue (not indicated) based on the provided molecular targets and references cited.

### Aire deficiency model

#### Human Aire-dependent genes

We followed previous studies [26, 105, 106] to obtain AIRE-dependent genes in human using orthologs of murine Aire-dependent genes [3]. Aire-dependent genes were defined by Sansom *et al.* (2014) as differentially expressed genes between *Aire* knock-out mTECs and mature *Aire* expressing mTECs in mice (Benjamini-Hochberg corrected pvalues ≤ 0.05). Supplemental Table 3, sheet 16 of Sansom *et al.* (2014) [3] provides all differentially expressed genes, and applying a fold change threshold of ≤ 2 yields the 3,980 AIRE dependent genes described in the paper. We matched this set of *Aire*-dependent genes to human orthologues, using *biomaRT* (v2.46.3 [107]), via attribute *hsapiens_homolog_ensembl_gene*, to obtain 3,361 unique human AIRE dependent genes (ensembl gene identifiers).

#### Generalization under AIRE-deficiency

To model negative selection in an AIRE-deficient thymus, we estimated thymus peptide abundances by removing the 3,361 human orthologs of Aire-dependent genes from the thymus bulk expression data, followed by peptide abundance estimation (Methods, *Peptide abundance*). We then applied our negative selection model on the AIRE-deficient thymus and analyzed all TCRs with survival probability > 0.98; we also analyzed TCRs that had a > 50% increased chance of survival in the AIRE deletion versus the baseline model (Table 1, column 𝑁_TCRs_). For each set of TCRs, we obtained the union of peptides in their cross-reactivity balls (Table 1, column 𝑁_peptides_). We then used a paired, one-sided t-test to determine if these sets of peptides were enriched in APS1 tissues (adrenal gland, liver, ovary, pancreas, skin, small intestine, thyroid, testis) versus non-APS1 tissues (bladder, blood, blood vessel, brain, breast, cervix uteri, colon, esophagus, fallopian tube, heart, kidney, lung, muscle, nerve, pituitary, prostate, salivary gland, spleen, stomach, uterus, vagina), measured at the sum of peptide weights across the respective tissues. To estimate empirical null distributions on these statistics, we randomly chose 𝑁_TCRs_ TCRs for each condition from the set 52.8 million self-reactive TCRs, and repeated the above analyses. For each random test, we drew 100 random sets of TCRs and computed the p-value associated with their t-test statistic. To obtain an empirical p-value, we counted the fraction of how often our observed p-value was smaller than the p-value across random sets, out of the total number of random sets.

## Author contributions

Conceptualization: HVM and SN; Theoretical and computational analysis: HVM, AB, SD, CK and SN; Experimental data generation and processing: HVM, SRC, YL, RKP; Writing - original draft: HVM and SN; Writing - review & editing: all authors; Supervision: HVM and SN.

## Data availability

Preprocessed data will be available on Zenodo upon publication.

## Code availability

Custom analysis code was written in either R (version ≥ 4.3.3) or Python (version ≥ 3.6.8). All analyses code is freely available on GitHub: https://github.com/meyer-lab-cshl/central-tolerance-generalization.

## Acknowledgments

We thank Vasilisa Kovaleva, David Pattinson, Yunxin Xie, and QED science for helpful feedback on the manuscript; Vasilisa Kovaleva for help in generating figure schematics; and Faviel Gonzalez-Galarza for providing frequency files from http://www.allelefrequencies.net.

## Protection of low-weight self peptides

We found no correlation between the thymic weight of a self peptide and the average weight of all other self peptides in its cross-reactivity ball. In addition, while individual self peptide weights span over 7 orders of magnitude, the average expression density in the cross-reactivity ball was much tighter, spanning only roughly 1 order of magnitude.

**Supplementary Figure 1.**
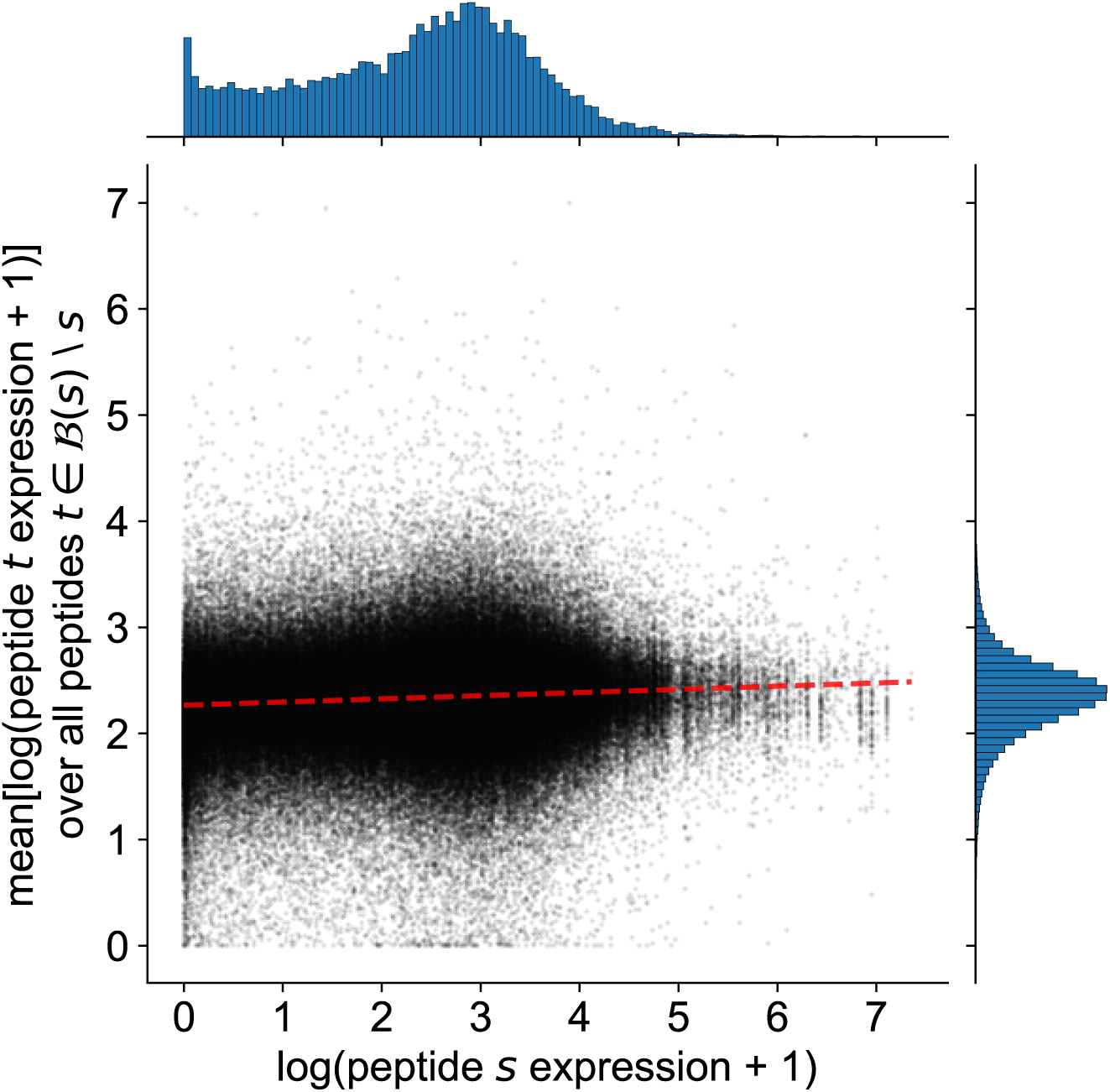
Single peptide expression versus cross-reactivity ball expression. The x-axis shows the expression of each self peptide 𝑠 in the thymus. The y-axis shows the average expression of all peptides in the cross-reactivity ball, *B*(𝑠) ⧵ 𝑠. Red dashed line shows Pearson correlation (𝑟 = 0.09).

## Generalization model applied to other MHC molecules

In humans, there are three highly polymorphic MHCI loci, which encode HLA-A, HLA-B and HLA-C molecules expressed simultaneously on the surface of the cell. We considered three additional molecules — A0101 from the A locus, B0801 from the B locus, and C0701 from the C locus — which each have a distinct profile of peptides bound. We applied our model to the union of MHC binders across these three loci, representing the spectrum of peptides that can sampled on an individual with the combination, or haplotype, of these loci. This haplotype is commonly found in four continental ancestries and is thus widely representative (Supplementary Fig 2). We found consistent results across the three individual loci, as well as the haplotype (Supplementary Table 1), i.e., sampling 10–20% of self peptides is sufficient to reduce peripheral damage by 85–90%.

**Supplementary Figure 2.**
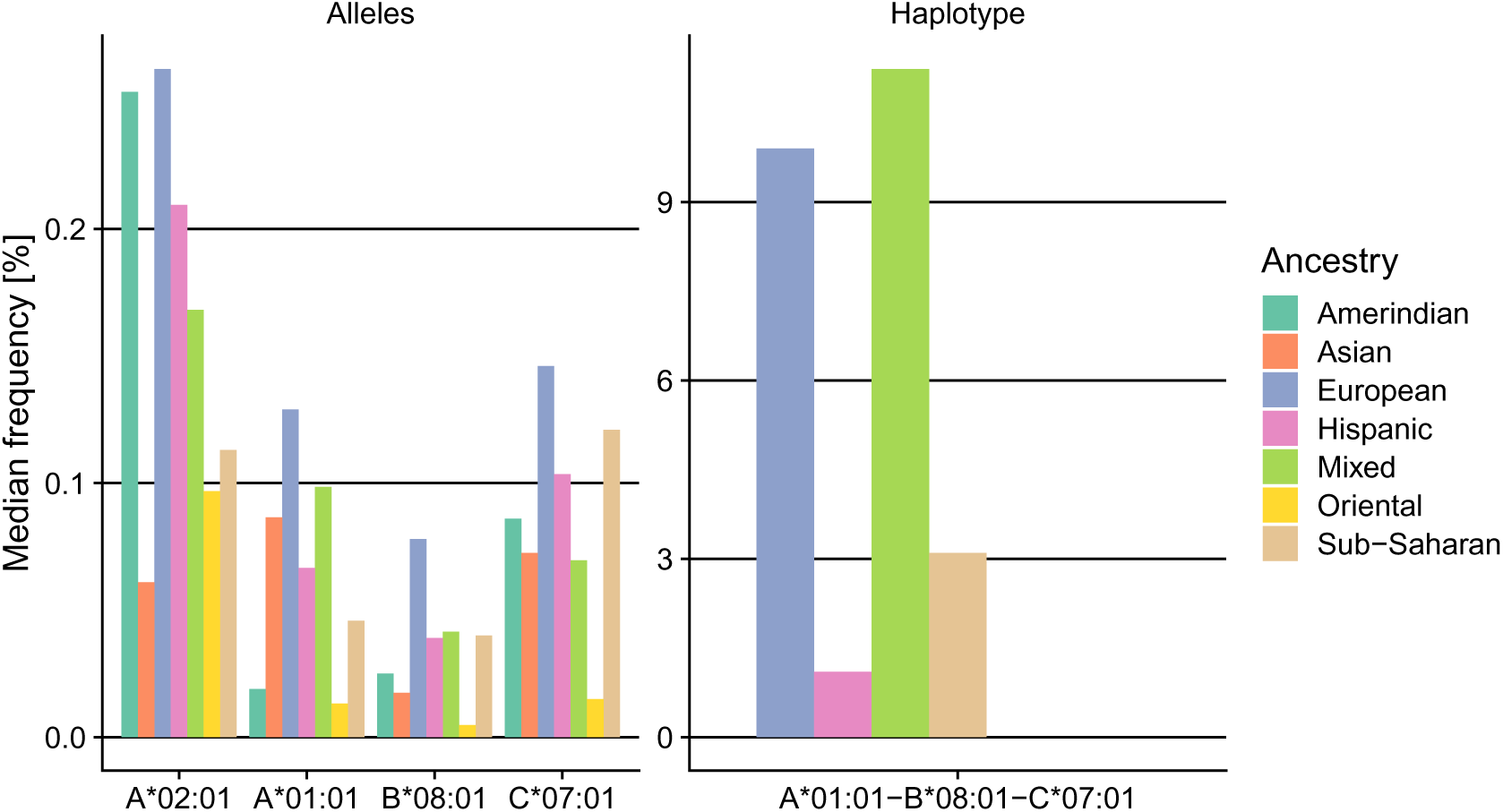
Haplotype frequencies. Haplotype frequencies at the MHCI locus for 7 broad continental ancestry groups. Frequency data from http://www.allelefrequencies.net [77].

**Supplementary Table 1.** Generalization model across MHC I loci. ‘Allele/Haplotype’ specifies the loci or their combination selected based on their high frequency across multiple populations (Supplementary Fig 2). ‘# MHC-binders’ is the number of 6mer binders derived as per Figure 1A. ‘# Seen’ shows the range of number of peptides sampled when varying the number of unique peptides per mTEC from 100 to 3,000. ‘% Protection’ was computed as 1 − Equation (3)∕Equation (1), and was estimated for a medium (100K) and a high (200k) estimate for number of peptides sampled.

| Allele/Haplotype | # MHC-binders | # Seen [Lower – Upper] | % Protection |  |
| --- | --- | --- | --- | --- |
|  |  |  | 100K samples | 200K samples |
| A0201 | 426,316 | 21,805 – 218,141 | 91.3 | 94.2 |
| A0101 | 306,861 | 21,614 – 178,524 | 87.7 | 91.5 |
| B0801 | 553,698 | 22,321 – 258,787 | 91.5 | 94.4 |
| C0701 | 703,380 | 22,706 – 293,600 | 89.2 | 92.9 |
| A0101-B0801-C0701 | 1,222,228 | 23,120 – 374,938 | 89.5 | 93.3 |

**Supplementary Figure 3.**
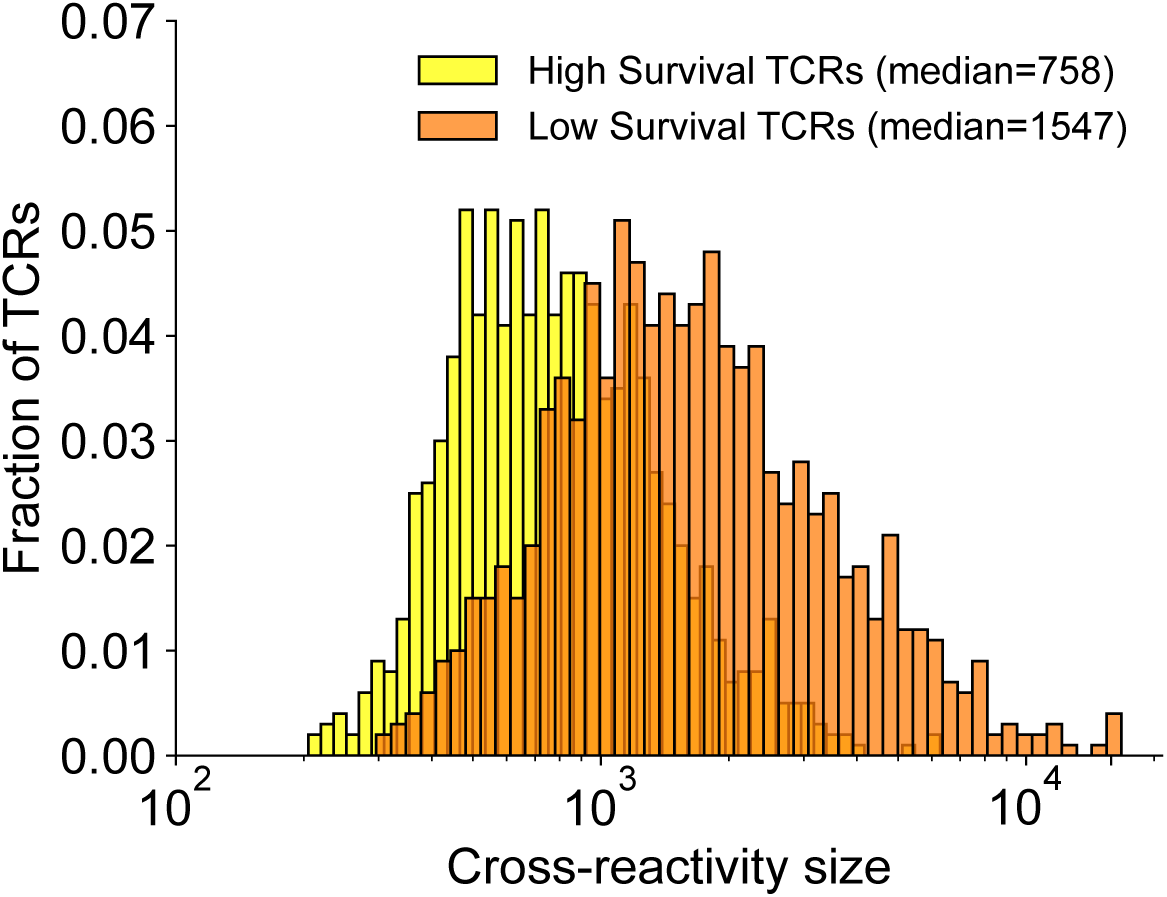
Negative selection narrows cross-reactivity. After applying our model (𝑘 = 100K), the number of total peptides in the cross-reactivity balls of ‘High Survival’ TCRs (defined as all TCRs 𝑡 with Pr_NS_(𝑡 survives) ≥ 0.5) is roughly 2-fold smaller than those of ‘Low Survival’ TCRs (Pr_NS_(𝑠 survives) < 0.5).

## Baseline generalization model vulnerabilities

We applied our negative selection model to the fully intact thymus and analyzed all TCRs with survival probability > 0.98, yielding 1,771,619 TCRs with a total of 53,694 peptides in their cross-reactivity balls. We then used a paired, one-sided t-test to determine if these sets of peptides were enriched in any of the 29 GTEx tissues compared the mean of the remaining 28 tissues. We adjusted this p-value according to Benjamini-Hochberg to account for the 29 statistical tests. To estimate empirical null distributions on these statistics, we randomly chose 100 sets of 𝑁_TCRs_ = 3900 TCRs from the set of 52.8 million self-reactive TCRs, which on average had a total of 53,700 self-peptides in their cross-reactivity balls, and repeated the above statistical analyses. To obtain an empirical p-value, we counted the fraction of how often our observed Benjamini-Hochberg adjusted p-value was smaller than the Benjamini-Hochberg adjusted p-value across random sets. Tissues with Benjamini-Hochberg adjusted p-value < 0.05 are indicated in gray outlines in Supplementary Fig 4 and their associated statistics are shown in Supplementary Table 2.

**Supplementary Figure 4.**
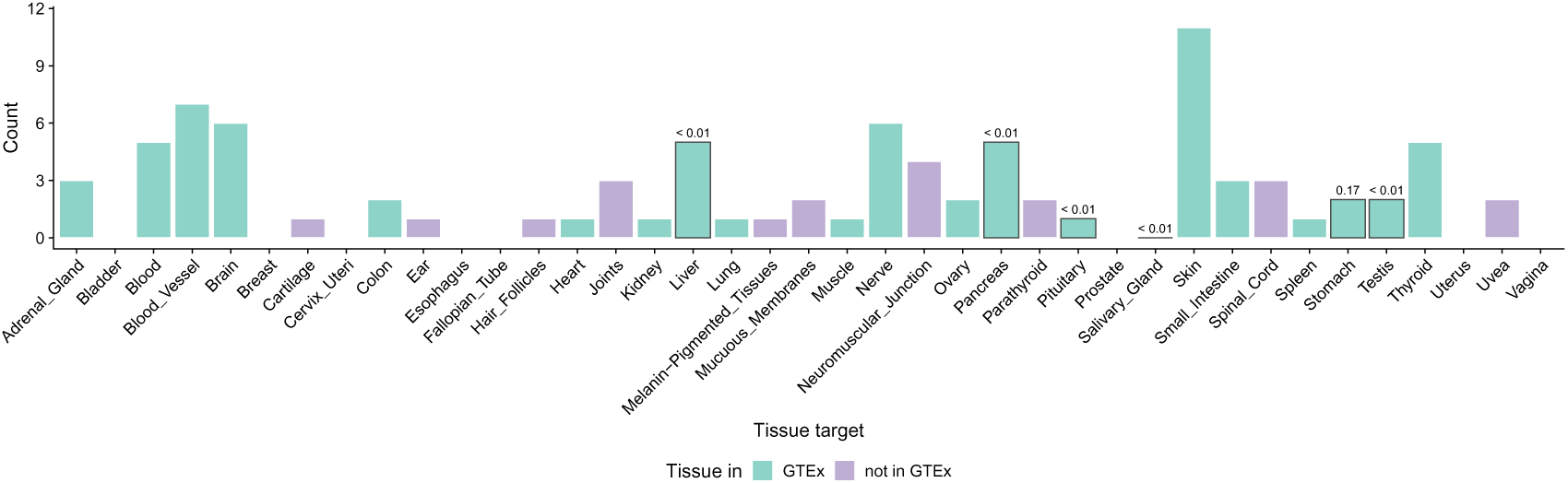
Immune disease statistics. Histogram of the number of times a tissue was identified as a target of the 81 autoimmune diseases defined by Hayter and Cook [104]. The color scale indicates if a tissue is present in the GTEx tissue collection. For each tissue in GTEx, we computed the enrichment of peptides within the cross-reactivity ball of TCRs with high survival probabilities (> 98%) using the baseline generalization model. Bars with gray outline indicate tissues with significant peptide enrichment, and values shown above the bar are empirical p-values in comparison to 100 random draws of TCRs and their corresponding target peptides.

**Supplementary Table 2.** Peptide abundance statistics in six tissues implicated in 15 autoimmune diseases. Summary of peptide enrichment statistics in six tissues with identified vulnerability in 15 autoimmune diseases (Supplementary Fig 4, gray outlines). ‘Mean abundance’ specifies the peptide abundance in the target tissue (first column) versus the other 28 GTEx tissues (second column) for peptides within the cross-reactivity ball of TCRs with high survival probabilities (> 98%) in the baseline generalization model. ‘p.adjust’ is the Benjamini-Hochberg adjusted ‘p-value’ (adjustment for 29 tissue enrichment tests). ‘p.emp’ is the empirical Benjamini-Hochberg adjusted p-value in comparison to 100 random draws of TCRs and their corresponding target peptides. ‘FC’ specifies the fold change of the mean peptide abundance in the target tissue over the other tissues.

| Tissue | p-value | p.adjust | p.emp | mean abundance |  | FC |
| --- | --- | --- | --- | --- | --- | --- |
|  |  |  |  | target tissue | other tissues |  |
| Liver | 3.66E-15 | 5.30E-14 | <0.01 | 11.37 | 6.88 | 1.65 |
| Pancreas | 3.17E-12 | 3.06E-11 | <0.01 | 48.97 | 5.53 | 8.85 |
| Pituitary | 0.009 | 0.044 | <0.01 | 16.21 | 6.70 | 2.42 |
| Salivary Gland | 5.47E-04 | 0.003 | <0.01 | 14.16 | 6.78 | 2.09 |
| Stomach | 7.83E-05 | 5.67E-04 | 0.17 | 20.74 | 6.54 | 3.17 |
| Testis | 1.13E-55 | 3.26E-54 | <0.01 | 12.72 | 6.83 | 1.86 |

